# Differential susceptibility of neuronal neurotransmitter phenotypes to HHV6 infection

**DOI:** 10.1101/2021.01.28.428686

**Authors:** E Bahramian, M Furr, JT Wu, RM Ceballos

## Abstract

Within the family *Herpesviridae*, sub-family *β*-*herpesvirinae*, and genus *Roseolovirus*, there are only three human herpesviruses that have been discovered and described: HHV-6A, HHV-6B, and HHV-7. Initially, HHV-6A and HHV-6B were considered to be simply two variants of the same virus (i.e., HHV6). Despite high overall genetic sequence identity (~90%), HHV-6A and HHV-6B are now recognized as two distinct viruses of the genus. Limited sequence identity (e.g., <70%) in key coding regions as well as significant differences in physiological and biochemical profiles (e.g., preferential use of different receptors for viral entry into susceptible hosts) underscore the conclusion that HHV-6A and HHV-6B are distinct virus species. Likewise, each virus appears to differentially contribute as putative etiologic agents to a variety of neurological disorders, including: multiple sclerosis, epilepsy, and chronic fatigue syndrome. Despite being implicated as causative agents in nervous system dysfunction, mechanisms of action and relative contributions of each virus to neural disorders remain elusive. Unresolved questions regarding: cell receptor use and binding affinity (i.e., CD49 versus CD134); cell tropism; the role of HHV-7 superinfection; and, relative virulence between HHV-6A versus HHV-6B – prevent a complete characterization. Although it has been demonstrated that both HHV-6A and HHV-6B can infect glia and, more recently, cerebellar Purkinje cells, cell tropism of HHV-6A versus HHV-6B for different nerve cell types remains vague. In this study, we demonstrate that both HHV-6A and HHV-6B can infect different nerve cell types (i.e., glia versus neurons) and different neuronal neurotransmitter phenotypes derived from the differentiation of human neural stem cells. We further show that both HHV-6A and HHV-6B induce cytopathic effects (CPEs) in susceptible nerve cells. However, the time-course and severity of CPEs appear to differ between HHV-6A versus HHV-6B infections and are dependent upon multiplicity of infection (MOI). As demonstrated by immunofluorescence, although both the HHV-6A and HHV-6B viruses productively infected VGluT1-containing cells (i.e., glutamatergic neurons) and dopamine-containing cells (i.e., dopaminergic neurons), neither HHV-6A nor HHV-6B challenge resulted in the productive infection of GAD67-containing cells (i.e., GABAergic cells). The reason underlying the apparent resistance of GABAergic cells to HHV-6A and HHV-6B infection remains unclear. Morphometric and image analyses of neurite extension and retraction dynamics as well as the time-course of cell aggregation phenomena (e.g., syncytia formation) during infection also indicate that HHV-6A induces more severe CPEs than HHV-6B at the same time-point and MOI. These data suggest that HHV-6A is more virulent than HHV-6B on susceptible human neural stem cells (HNSCs) differentiated into neuronal phenotypes, while neither virus is able to infect GABAergic cells. If these *in vitro* data hold *in vivo*, the *inhibitory interneuron dysfunction hypothesis* for HHV6-driven seizures may be ruled out as a potential mechanism for HHV6-induced epileptogenesis.

## 1. Introduction

Human herpesvirus 6 (HHV-6) variants fall into two sub-groups of viruses, which are now distinguished by the International Committee on Taxonomy of Viruses as distinct virus species, designated as HHV-6A and HHV-6B, in the genus *Roseolovirus* (Adams and Carstens, 2012). Along with human herpesvirus 7 (HHV-7), HHV-6A and HHV-6B are the only characterized human viruses included within the genus *Roseolavirus* (subfamily *β*-*herpesvirinae*, family *Herpesviridae*). Despite exhibiting approximately 90% overall genome sequence identity (Isegawa et al., 1999), key regions of the HHV-6A and HHV-6B genomes (i.e., immediate-early genes) exhibit only 70% (or less) sequence identity (Dominguez et al., 1999). Beyond notable gene sequence divergence, key coding regions yield homologous proteins with distinguishable amino acid profiles even when the genes exhibit high (e.g., 95%) sequence homology (Rapp et al., 2000; Achour et al, 2008). For example, HHV-6A and HHV-6B envelope glycoproteins (i.e., gH and gB), which are critical for virus-cell surface interactions, exhibit overall high sequence identity between homologous genes but feature distinct amino acid profiles for each of the two species (Achour et al., 2008). This may account for noted variations in cell tropism (and etiology) between the two viruses (Oyaizu et al., 2012; Tang and Mori, 2018). Until 2013, an inhibitory complement receptor, CD46 (cluster of differentiation 46) was considered to be the primary target for HHV-6A attachment to susceptible cell types (Santoro et al., 1999; Santoro et al., 2003). However, it has been suggested that tumor necrosis factor receptor superfamily member 4 (TNFRSF4), also known as CD134, serves as the primary receptor for HHV-6B entry into susceptible cell types (Tang et al., 2013). Indeed, there may be other yet to be identified cell surface receptors to which HHV-6 virions can bind with asymmetric binding affinity between such receptors and HHV-6A versus HHV-6B envelope glycoproteins. This area of research is ongoing.

Among the tissues and organs that are known to harbor HHV-6A and HHV-6B, it is known that both viruses can infect the central nervous system. (De Bolle et al., 2005; Boutolleau et al., 2006). Although reports suggest that HHV-6A may be more neurotropic than HHV-6B (Hall et al., 1998), this is mainly based on the prevalence of HHV-6A over HHV-6B in cerebral spinal fluid and blood of patients with rhomboencephalitis, multiple sclerosis, and other neuroinflammatory diseases (Crawford et al., 2007; Alvarez-Lafuente et al., 2010). In general, there is a lack of evidence showing the extent of HHV-6A versus HHV-6B cell tropism in the central nervous system (CNS). It is unknown whether the brain proper or nerve cells within the spinal column exhibit predisposed susceptibility to one virus over the other. In the brain, it is unknown if there is predisposition for HHV-6A versus HHV-6B infection in glial versus neurons. Once it became clear that HHV-6A and HHV-6B were two distinct viruses, it was demonstrated that both were able to infect astrocytes (Fotheringham et al., 2008). Since then, a few studies have emerged differentiating between HHV-6A and HHV-6B infection in select nerve cell types (Prusty et al., 2018; Liu et al., 2018). However, such studies are limited (*see review* Santpere et al., 2020). The relative virulence of HHV-6A versus HHV-6B on susceptible nerve cell types and differential susceptibility of specific neuronal neurotransmitter phenotypes to these two viruses remains unclear. Characterization of infection dynamics and cell tropism of HHV-6A versus HHV-6B is essential for validating models of HHV6-based neurological dysfunction. Recently, a study was published demonstrating that HHV-6A and HHV-6B infect Purkinje cells (Prusty et al., 2018). Given that Purkinje cells are GABAergic, the data from that study struck our interest since we were unable to detect the presence of HHV-6A or HHV-6B in GABAergic differentiated human neural stems cells (HNSCs). In this study, we provide data from an immunofluorescence (and qPCR) study confirming that both HHV-6A and HHV-6B infect both GFAP-positive cells (i.e., glia) and βIII-tubulin-positive cells (i.e., neurons) with notable cytopathic effects. We also show that both viruses can infect different neuronal neurotransmitter phenotypes. However, neither HHV-6A nor HHV-6B appears to infect GABAergic cells. In cells which are susceptible, the timing of onset for cytopathic effects (CPEs) and CPE severity differs between HHV-6A versus HHV-6B infections under equivalent conditions.

## 2. Materials and methods

### 2.1 Cell culture

Culture vessels (i.e., T75 flasks and 8-well microscope chamber slides; Thermofisher Scientific) were coated with CELLStart™ surface substrate (Gibco, Life Technologies Corporation) per supplier protocol. In serum-free medium (SFM) [Knockout™ DMEM/F-12, 20 ng/mL FGF-2/EGF, 2mM GlutaMAX™-I, and 2% v/v StemPro™ Neural supplement], NIH-approved H9-derived human embryonic stem cells (hESCs) were plated and expanded as a monolayer in CELLStart™ coated vessels. The cultures were maintained at 37°C in a humidified incubator (with 5% CO_2_). Cells were passaged every 7 days using an accutase cell detachment solution (Sigma-Aldrich) and re-plating at a surface density of 5 × 10^4^ cells/cm^2^. Using an automated cell counter (Bio-Rad) and Trypan Blue (Thermofisher Scientific), viable cell counts were determined for re-plating cells. Upon re-plating, different media options were used to induce differentiation along desired paths. For example, to facilitate differentiation toward neuron-dense mixed cultures, a seeding density of 2.5 × 10^4^ cells/cm^2^ was used with a minimal media (MM) [Knockout™ DMEM/F-12, 2mM GlutaMAX™-I, and 2% StemPro™ Neural Supplement]. Cells were maintained in differentiating conditions for 15-18 days with fresh media change-out every 3-4 days.

### 2.2 Virus preparation

Frozen stocks (−195°C) of HHV-6A strain GS-infected HSB2 cells and HHV-6B strain Z29-infected MOLT-3 cells (courtesy NIH) were thawed and used to infect uninfected HSB2 and MOLT-3 cells. Specifically, 10^6^ HHV6-infected cells were mixed with uninfected cells at a ratio of 1:10 in a T150 flask (Thermofisher Scientific) and incubated at 37°C for 2 hr. (5 mL of media). After 2 hr of incubation fresh media as added (5 mL) and the cells were incubated again at 37°C. Using a light microscope, the culture was checked periodically. When cytopathic effects (CPEs) were noted in more than 80% of cells, the cell suspension was harvested. Aliquots were stored in liquid nitrogen (−195°C) for use in infection assays. Alternatively, cell-free virus suspension was prepared by sonicating cell suspension on ice and centrifuging the lysate at 3500 rpm for 1 hr to pellet cell debris while maintaining virus particles in suspension. The supernatant was extracted and filtered through a 0.45μm filter to remove any remaining cell debris. The virus-containing filtrate was centrifuged at 25,000 rpm at 4°C for 3 hr to pellet the virus. Supernatant was removed and the virus pellet was resuspended in cold media and stored at −80°C for use in cell-free virus infection assays or for transmission electron microscopy. Virus titers were determined using qPCR.

### 2.3 Transmission electron microscopy (TEM) for cell-free virus

Transmission electron microscopy (TEM) was used for three purposes. First, TEM was used to verify the presence of intact fully-assembled virions from HSB2 and MOLT-3 host cells used for virus storage and propagation. Second, TEM was used to demonstrate the presence of HHV6 virus particles in differentiated HNSCs. Third, TEM was used to validate the presence of virus for qPCR-based titers from storage cells or differentiated HNSCs. For cell-free virus samples, the aforementioned preparation via sonication (or, alternatively, freeze-thaw cycles) was followed by concentration of virus suspensions (i.e., filtered lysates) using 30kDA MWCO spin concentrator (Sigma-Millipore). From the retentate, ~5μL of concentrated viral suspension was spotted onto a formvar-coated copper grid and incubated for 10 min in a humidity chamber. The sample was then gently rinsed and negatively-stained with a 2% solution of uranyl acetate for 30 sec before excess solution was wicked off of the grid and allowed to air dry for 1 hr. Samples placed on grids, and after 2 to 10 minutes, 2% uranyl acetate was added to the grid. Grids were imaged with a Hitachi H-7100 TEM at 75 kV. Images were captured at 60,000– 200,000X magnification.

### 2.4 Transmission electron microscopy (TEM) for infected cells

TEM was also used to image virus particles associated with host cells. These included both virus storage cells (HSB2 and MOLT-3 cells) and differentiated HNSCs infected with either HHV6 virus. For virus-infected storage cells in or for virus-infected monolayers of differentiated HNSCs, cells were fixed with a 4% paraformaldehyde (PFA) solution. The samples were place on grids and then gently rinsed and negatively-stained with a 2% solution of uranyl acetate. Samples were incubated for 2 hr at 4°C before rinsing with distilled water and wicking off excess solution from the grid and allowed to air dry for 1 hr. After air drying for 2 hr., samples were imaged with a Hitachi H-7100 TEM at 75 kV. Images were captured at 60,000– 200,000X magnification. Regions of the grid that showed virions blebbing from membrane or virus particles within the intracellular space were targeted for imaging. Detachment of HNSCs from the surface substrate was required, to capture co-localization of virus particles within intact cells.

### 2.5 Light microscopy

The infected cultures were monitored daily via light microscopy to monitor morphological changes. (In each experiment, uninfected cultures served as control). Light microscopy images were taken from dHNSCs at PDD 7 and PDD 14 at 2 hours post-infection (HPI) and 24 HPI10 using an upright microscope (Primovert, Zeiss) equipped with a color camera (AxioCam 105, Zeiss). Dozens of cells were captured in each image from randomly chosen fields of view for each culture.

### 2.6 Morphometric Analyses

Using light microscopy and an image analysis pipeline that includes a machine learning algorithm to enhance calls on neurite extension, node formation, and other morphometric parameters, cell morphology changes are monitored over the course of infection and at different infection states. Using the NeuronJ plug-in to the ImageJ software package and a machine-learning algorithm, initial manual tracing of neurons from light microscopy images can be enhanced. This first part of the workflow compares pixel intensity on neurites (and soma boundaries) with adjacent pixel neighborhoods. Using the machine-learning algorithm, Trainabler Weka Segmentation 7, the contrast between nerve cell features and background is enhanced yielding *pseudo-fluorescence* images from light microscopy images. Standard tools such as Neural Circuit Tracer or NeuronCytoII are then employed to generate reconstructions of nerve cells allowing for morphometric analysis without fixing cells for traditional immunofluorescence. This workflow requires minimal high-performance computing time and allows observations and quantification of morphological parameters in real time while producing quality images (*see* Fig. 6, panels A-C).

### 2.7 Immunofluorescence and fluorescence microscopy

Immunofluorescence was conducted via a co-labeling approach (two antibody systems per trial) along with the nuclear dye 4,6-diamidino2-phenylindole dihydrochloride (DAPI). The following fluorescence antibody systems (primary antibody/secondary antibody) were used in select pairs: mouse anti-gB (HHV6 envelope glycoprotein gB)/donkey anti-mouse IgG-Alexa Fluor 488; chicken anti-GFAP (Glial Fibrillary Acid Protein)/donkey anti-chicken IgG-Alexa Fluor 680; rabbit anti-βIII tubulin (neuron-specific microtubule protein)/donkey anti-rabbit IgG-Alexa Fluor 568; goat anti-VGluT (vesicular glutamate transporter protein)/donkey anti-goat IgG-Alexa Fluor 555; chicken anti-GAD67 (glutamate decarboxylase 67)/donkey anti-chicken IgG-Alexa Fluor 680; and, rabbit anti-DA (dopamine)/donkey anti-rabbit IgG-Alexa Fluor 568. The anti-gB antibody (courtesy NIH AIDS reagent program) and fluorescent secondary indicate the presence of HHV6 virus and when co-localized with other immunofluorescence markers demonstrate infection in select cell types (i.e., neurons versus glia or distinct neuronal neurotransmitter phenotypes).

Signal for anti-gB is usually color-coded green in alignment with emissions during image analysis. Signal for the anti-βIII tubulin fluorescent antibody system (Sigma-Aldrich) is typically color-coded red in alignment with emissions when anti-gB is co-labeled but may also be color-coded green in uninfected controls. Signal for the anti-GFAP fluorescent antibody system (Sigma-Aldrich) is typically color-coded red in alignment with emissions but may also be color-coded green in uninfected controls or when used as a co-label for distinguishing between GFAP-positive cells and βIII tubulin-positive cells in the same image. Signal for anti-VGluT (Abcam), anti-GAD67 (Abcam), and anti-DA (EMD Millipore) fluorescence is typically color-coded red to distinguish neuronal neurotransmitter phenotype from any anti-gB signal (green). All experimental trials used DAPI (blue color code) to locate nuclei of all cells in the mixed cultures regardless of cell type. For all immunofluorescence trials, HNSCs were fixed with 4% PFA in Dulbecco’s phosphate buffer solutions (DPBS), then rinsed (3X) with DPBS. Blocking solution (5% donkey serum, 0.1% triton in DPBS) was added for 30 min. at room temperature (RT) after which primary antibodies (abs) were applied. Cells were then incubated for 24 hrs. followed by a series of rinses (3X) with DPBS and then application of secondary antibodies and further incubation. Another series of rinses (3X) with DPBS was followed by application of DAPI (0.2 nM) and a 20 min incubation at RT. The plates were subjected to a final rinse in DPBS and staged within the confocal fluorescent microscope (Leica). Image were taken using 10X magnification to view the distribution of fluorescent signals across the plates. In infecdted cultures, immunofluorescence was performed at multiple time-points during the differentiation phase (PDD7 and PDD14 preferred) and at several time-points after introduction of viral inocula (e.g., 2HPI and 24HPI). Images for each fluorescent signal were captured and then images were overlayed using image analysis software to generate composite images.

### 2.8 Quantitative polymerase chain reaction (qPCR)

For quantitative polymerase chain reaction (qPCR)-based virus titers from both storage cell lines (i.e., HSB2 and MOLT-3 cells) as well as infected differentiated HNSCs, the HHV6-specific U22 gene was targeted. Using the following primers (Collot et al., 2002): 397F (5’-TCG AAA TAA GCA TTA ATA GGC ACA CT-3’) and 493R (5’-CGG AGT TAA GGC ATT GGT TGA-3’) – a 99bp fragment of the U22 gene was amplified from viral DNA extracted from infected cell cultures. Amplification was performed using a Rotor-Gene Q real-time fluorescence detector thermocycler (Qiagen) programmed as follows: 94°C for 6 min; followed by 40 cycles of 94°C for 30 sec, 53°C for 30 sec, 72°C for 45 sec; and, then a final holding condition of 72°C for 7 min. Verification of the appropriate fragment size (and run quality) was checked via gel electrophoresis. DNA concentration was determined using a micro-volume spectrophotometer (Denovix). The number of viral genomes for each sample was determined by comparing results to a standard curve.

To prepare standards for qPCR analyses, the following procedure was performed for each virus (i.e., HHV-6A and HHV-6B): one copy of the U22 target sequence was cloned into a commercial vector using a TOPO PCR Cloning Kit (Invitrogen) and transformed into competent *E. coli*. Enrichment of clones was followed by plasmid purification using the QIAprep Spin Miniprep Kit (Qiagen). The resulting plasmid DNA yield was measured by absorbance spectroscopy (OD_260nm_). PCR is then employed to amplify the gene of interest (i.e., U22) from the plasmid preparation. PCR product yield is quantified using the Denovix micro-volume spectrophotometer and then a sample is diluted down to 1 ng/μL which corresponds to a fragment copy number of 9.216 × 10^9^. From this, subsequent dilutions (10^−1^ through 10^−9^) are prepared in triplicate. Next, qPCR is used to amplify fragments from these serial dilutions and the standard curve is generated. For all qPCR trials, 10μL SYBR^®^ Green (Life Technologies, Foster City, USA), 1 μL of unknown DNA sample, 50 nM of forward primer, and 50 nM of reverse primer was used in 20 μL total reaction volumes.

## 3. Results

### 3.1 Both glia and neurons are susceptible to infection by either HHV-6A or HHV-6B

Results from immunofluorescence histochemistry indicate that that both HHV-6A and HHV-6B can infect glial fibrillary acidic protein (GFAP)-positive differentiated human neural stem cells (HNSCs). Labeling HNSC cultures (H-9 derived cells, Gibco™) at post-differentiation day 7 (PDD7) with antibodies against GFAP and HHV6 envelope glycoprotein gB and co-staining with DAPI (4’,6-diamidino-2-phenylindole), reveals co-localization of anti-gB and anti-GFAP signals in DAPI-stained cells (Fig. 1). This indicates the susceptibility of glial cells to infection by HHV-6A (Fig. 1, row A) and HHV-6B (Fig. 1, row B). These data also show that by 2 HPI, cell aggregation, a common cytopathic effect (CPE) has begun. Distribution of cells in the uninfected control state (Fig. 1, row C) is notably more homogenous.

**Fig. 1.**
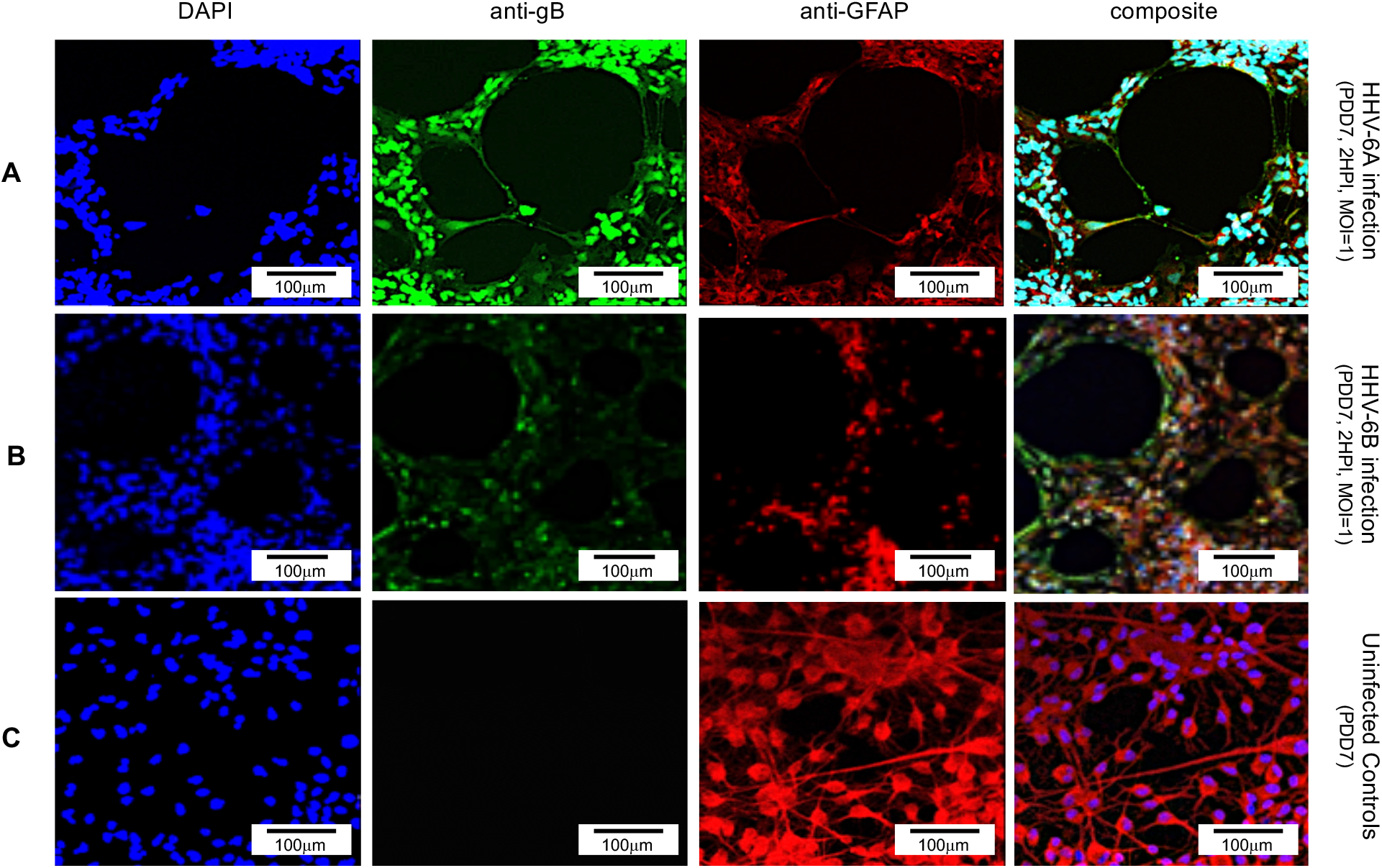
Fluorescence microscopy images of differentiated human neural stems cells (dHNSCs) treated with immunofluorescent antibodies and a fluoro-dye at PDD7: Differentiation to Glial cells. From left to right, DAPI, anti-gB, anti-GFAP, and composites. PDD7 HNSCs infected with HHV-6A (*row A*) show gB-positive signal on GFAP-positive cells (glial cells). PDD7 HNSCs infected with HHV-6B (*row B*) also show gB-positive signal on GFAP-positive cells. Images for uninfected control culture (row C) show well-developed GFAP-positive cells with homogeneous distribution; many exhibit stellate morphotypes.

Using an antibody against βIII-tubulin (a neuron-specific microtubule protein) in addition to DAPI and an anti-gB fluoroprobe, immunofluorescence demonstrates that neurons are also susceptible to infection by HHV-6A (Fig. 2, row A) and HHV-6B (Fig. 2, row B). Fluorescence images from the βIII-tubulin antibody system in the uninfected control trial (Fig. 2, row C) show elongated neurite extension and node formation further indicating HNSC differentiation into neurons and the formation of cell-cell connectivity. Reductions in cell density (i.e., cell death) and aggregation of infected cells after 2 HPI in PPD7 cells illustrate viral-induce CPEs in cells infected with HHV-6A (Fig. 2, row A) and HHV-6B (Fig. 2, row B).

**Fig. 2.**
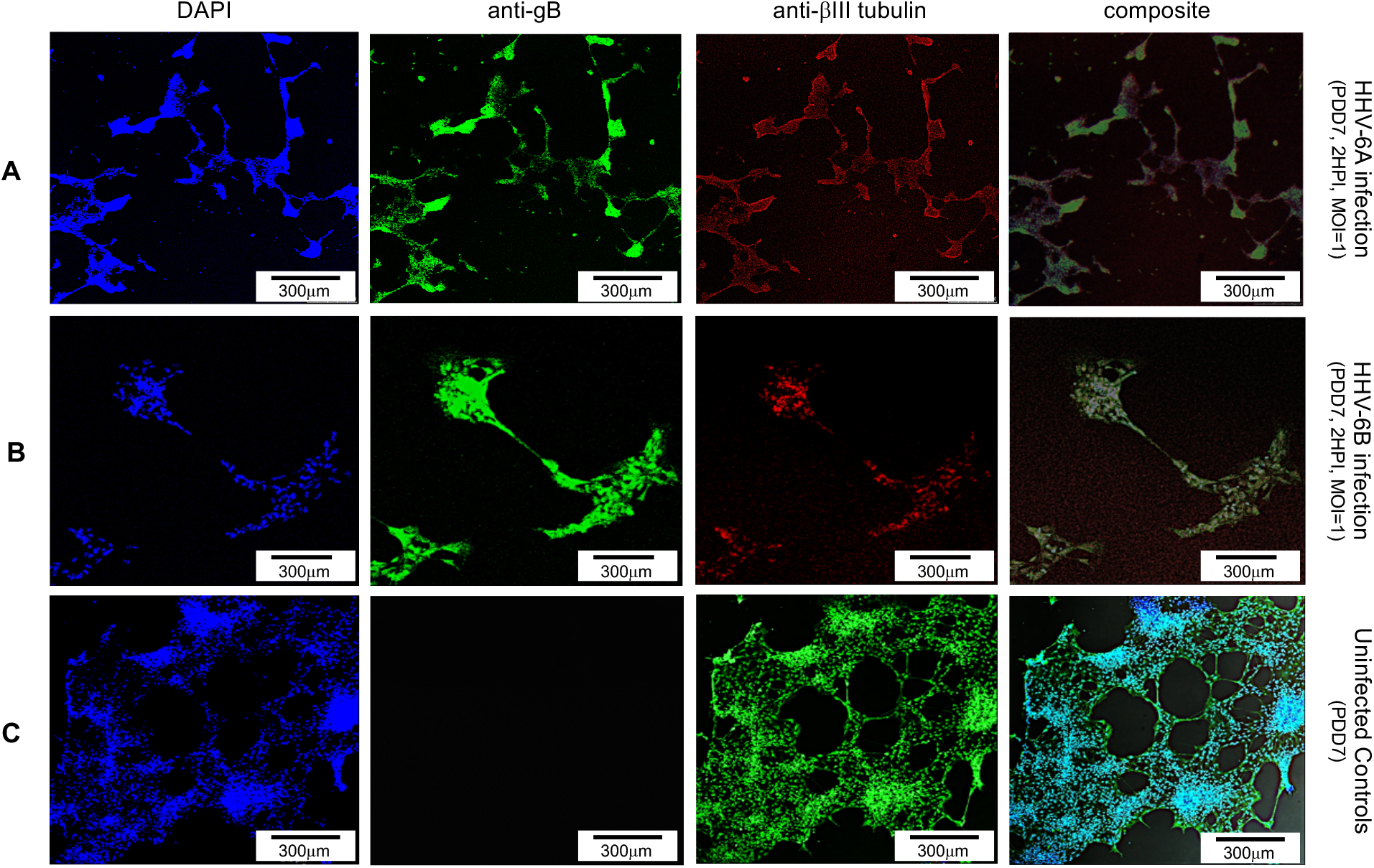
Fluorescence microscopy images of differentiated human neural stems cells (dHNSCs) treated with immunofluorescent antibodies and a fluoro-dye at PDD7: Differentiation to Neurons. From left to right, DAPI, anti-gB, anti-βIII tubulin, and composites. PDD7 dHNSCs infected with HHV-6A (*row A*) show gB-positive signal on βIII tubulin-positive cells. PDD7 HNSCs infected with HHV-6B (*row B*) also show gB-positive signal on βIII tubulin-positive cells. Images for uninfected control culture (*row C*) show developed βIII tubulin-positive cells with significant neurite-neurite and neurite-soma connectivity.

### 3.2 Infection with either HHV-6A or HHV-6B results in time-dependent cytopathic effects

Productive infection can be verified via transmission electron microscopy (TEM) and quantitative polymerase chain reaction (qPCR). TEM demonstrates the presence of fully assembled HHV-6A virus particles (Fig. 3A) within cells that match the approximate size and morphology of cell-free virions from virus stocks (Fig. 3B). Likewise, HHV-6B virus particles can be observed within a vacuole-like space within HNSCs (Fig. 3D). These also match the approximate size and shape of HHV-6B virions imaged from virus stocks (Fig. 3E). To further demonstrate that these are productive infection as opposed to a *lysis-from-without* phenomenon (Delbruck et al., 1940), qPCR-based virus titers (i.e., number of viral genomes per milliliter) were determined for HHV-6A (Fig. 3C) and HHV-6B (Fig. 3F). Within a couple of hours post-infection, the number of detectable viral genomes exceeds the virus density of the inoculum of HHV-6A and HHV-6B used to infect HNSC cultures, thereby demonstrating productive infection (i.e., the production of progeny virus).

**Fig. 3.**
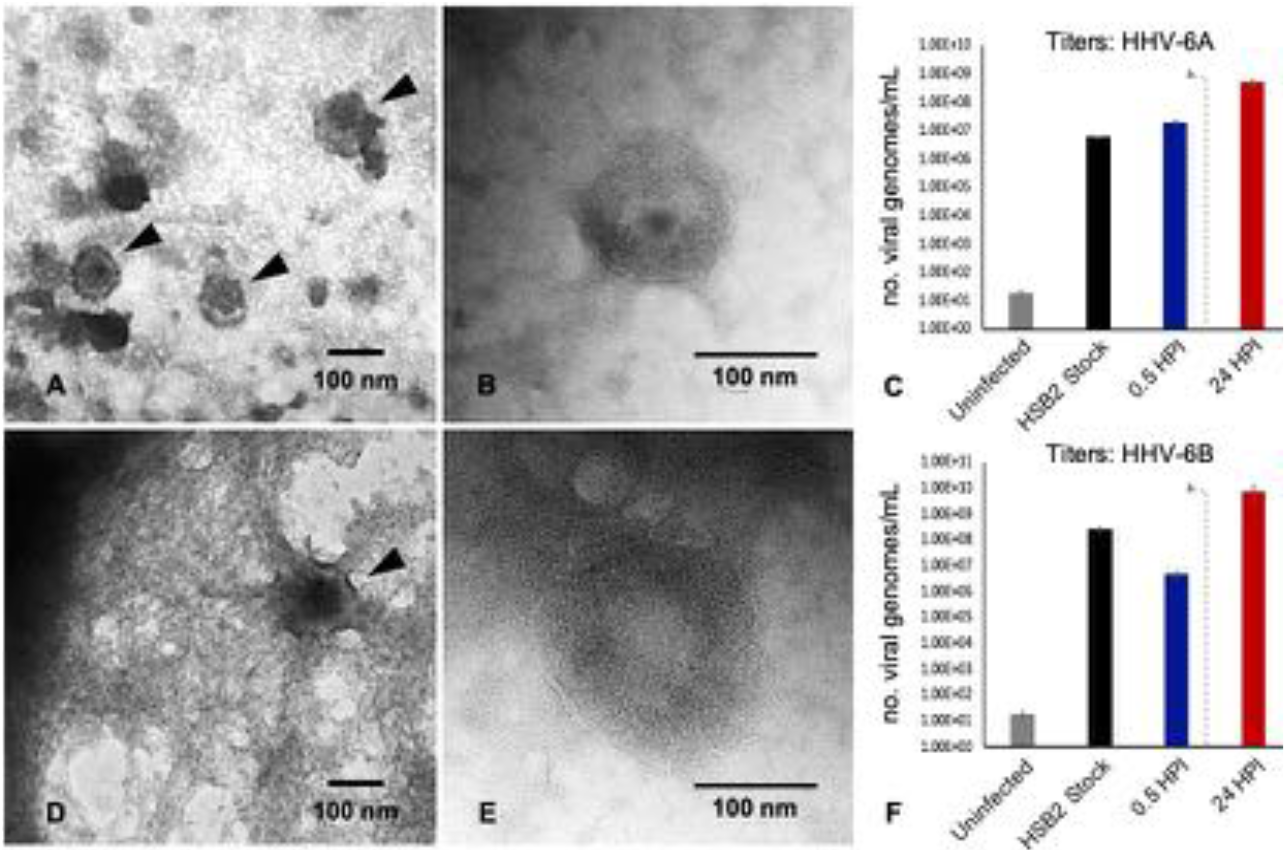
TEM and qPCR data from HHV6-infected hNSCs. TEM images illustrate: (A) HHV-6A virus particles within a cell; (B) HHV-6A virions in cell-free filtered supernatant from cell lysate; (D) HHV-6B virions in a cell; (E) HHV-6B virions in cell-free filtered supernatant. qPCR titer methods show productive virus infection for: (C) HHV-6A and (F) HHV-6B. Uninfected cells show negligible amplification of an HHV6-specific marker (i.e., U22 gene). After 30 min post-infection HHV6 is still present. After a wash at 2 HPI (dashed line) and 22 h incubation (24HPI), titers increase indicating production of progeny virions at densities greater than present immediately after inoculation.

Together, data from TEM, immunofluorescence using an anti-gB fluorescent antibody system, and qPCR-based quantification of virus titers at different time-points during infection, provide convincing evidence that both neurons and glia are susceptible to HHV-6A and HHV-6B and result in productive viral infections. Furthermore, both TEM and immunofluorescence data indicate that infection of dHNSCs by either HHV-6A or HHV-6B results in viral-induced CPEs, which manifest as disturbances to cell shape, size, viability, and distribution on culture plates. CPEs often present in two phases: cell aggregation (with notable cell death) and detachment from the culture surface.

#### 3.2.1. Severity and timing of CPEs differs between HHV-6A versus HHV-6B infection

The severity and time-course of CPEs differs between HHV-6A and HHV-6B infections (Fig. 4). Although there is notable aggregation of cells infected with either HHV-6A or HHV-6B after 2HPI at MOI = 1 or 2, HHV-6A infected cells produce higher density clumps at the earlier time point for both PDD7 and PDD14 cultures (Fig. 4A and 4C, *middle row*). By 24 HPI, HHV-6A infections yield high density detached clumps floating in culture (Fig. 4A and 4C, *bottom row*). Although HHV-6B infections induce cell aggregation under equivalent conditions, cells are more resistant to forming high-density spherical clumps that detach from the surface at 2 HPI and 24 HPI (Fig. 4B and 4D).

**Fig. 4.**
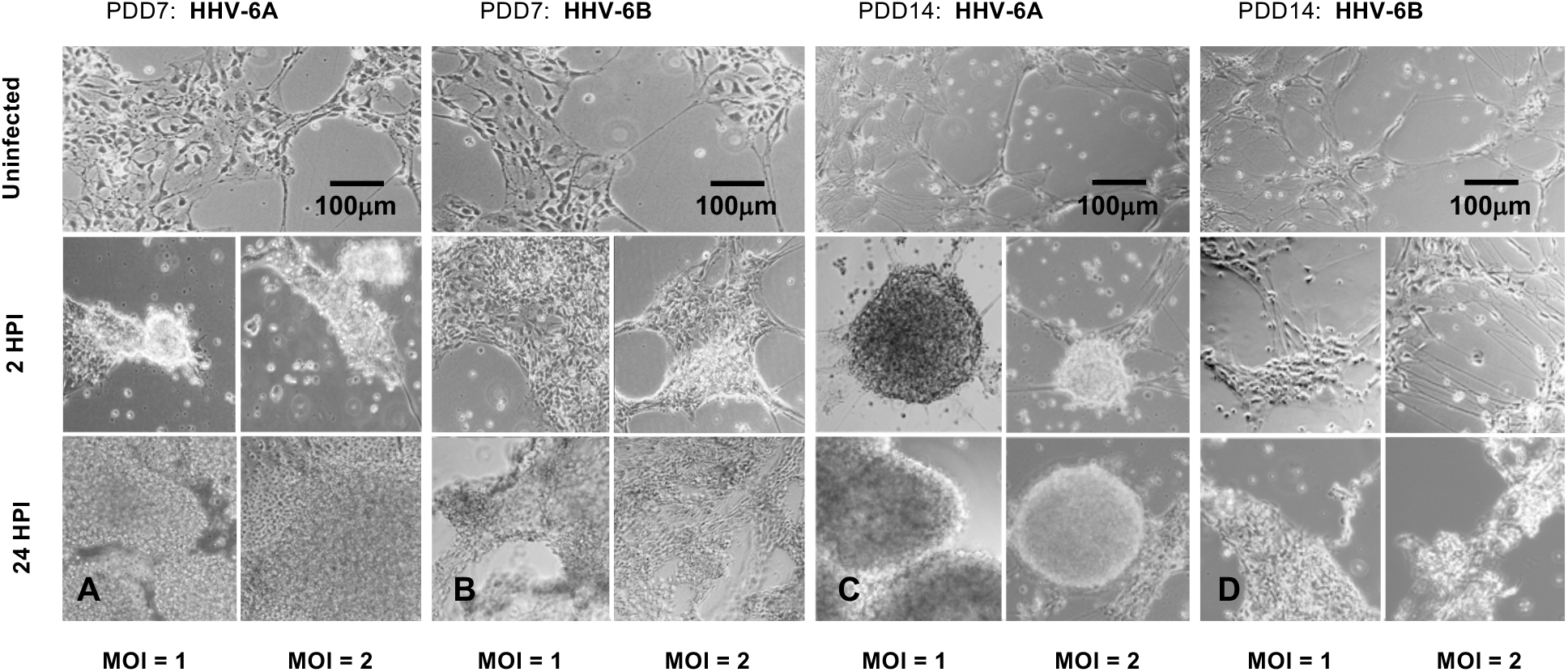
Light microscopy of HHV6-induced cytopathic effects (CPEs) on differentiated HNSCs at PDD 7 and 14. (A) Uninfected cultures of HNSCs at PDD7 show healthy cells adhering to the plate surface (*top*). After two hours post-infection (2HPI) at MOIs = 1, 2 (*middle, left and right, respectively*) with HHV-6A, cells begin to aggregate and at 24HPI with HHV-6A at MOIs = 1, 2 (*bottom, left and right*) there is high-density clumping and cell detachment. (B) Uninfected cultures of HNSCs at PDD7 show healthy cells adhering to the plate surface (*top*). After 2HPI with HHV-6B at MOIs = 1, 2 (*middle, left and right*), cells begin to aggregate and at 24HPI with HHV-6B at MOI = 1, 2 (*bottom, left and right*) there is higher-density aggregation. (C) At PDD14, uninfected healthy cells persist (*top*); however, at 2HPI at MOIs = 1,2 (*middle, left and right*) infection with HHV-6A results in high-density cell aggregation and at 24HPI rampant cell death and detachment is observed (*bottom, left and right*). (D) At PDD14, uninfected cells persist (*top*); however, HHV-6B infection at MOIs = 1,2 for 2 HPI results in lower-density cell aggregation (*middle, left and right*) with higher-density clumping occurring at 24HPI (*bottom, left and right*).

Indeed, in some trials, HHV-6B infected cells retained morphological integrity at 2HPI (Fig. 4D). To determine whether clumping of cellular biomass during the course of HHV6 infection is simply aggregation or *bona fide* viral-induced sycyntia formation, fluorescence microscopy was used to detect morphological features characteristic of syncytia (Fig. 5). Results indicate that prior to gross detachment from the culture surface (i.e., 0-2 HPI), morphological features consistent with syncytia occur, including cell fusion and formation of multi-nucleated larger cells (Fig. 5A; *arrows*). This occurs in both HHV-6A and HHV-6B infections. However, in uninfected control cultures, individual cells with robust arborized neurites and a single nucleus will persist for weeks (Fig. 5B). In uninfected cultures, viable nerve cells also feature slightly larger, well-defined soma boundaries and baseline electrical activity (not shown). (Details regarding changes in electric activity in HNSC cultures are part of a separate report).

**Fig. 5.**
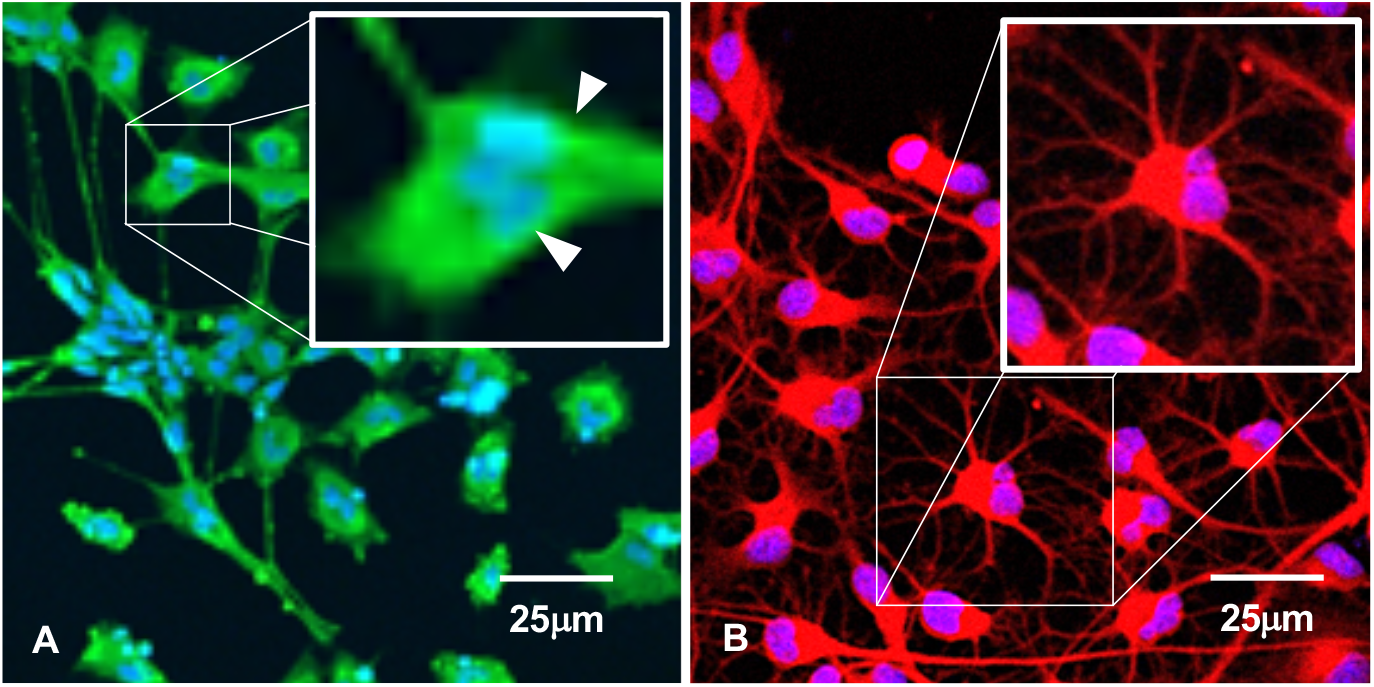
Immunofluorescence suggests syncytia formation. (A) HHV-6B infection in HNSCs results in syncytia formation as indicated by cell membrane fusion and multinucleated cells (arrows). (B) Uninfected culture shows individual well-bounded dHNSC membranes and the absence of any cell aggregation that would suggest syncytia-like formations.

**Fig. 6.**
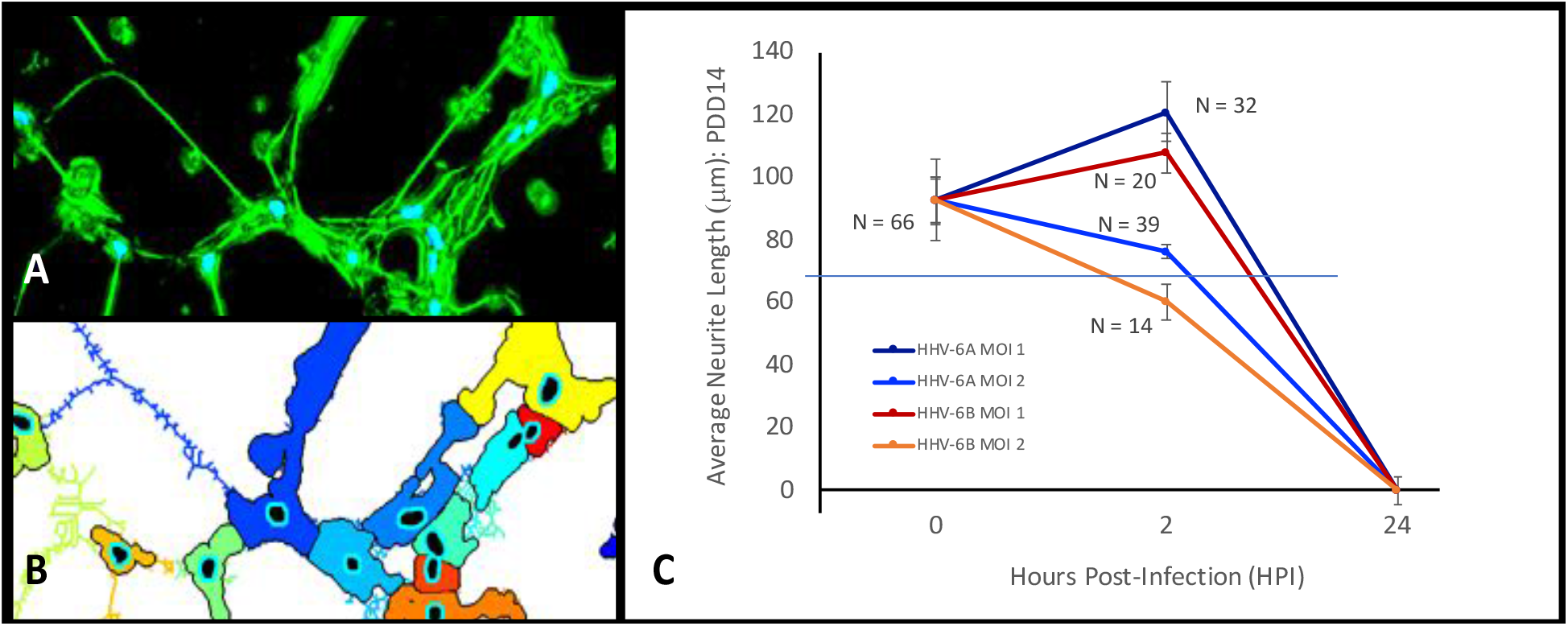
Changes in average neurite length and neuronal signaling during the course of an HHV6 infection. (A) Pseudo-flourescence image generated by Trainabler Weka Segmentation 7 highlights soma boundaries and neurites of neurons from light microscopy images processed via a NeuronJ plug-in to the ImageJ software package. (B) Neuron CytoII software generates neuron and neurite reconstructions for detailed morphometric measurements. (C) Morphometric analysis of average neurite length (ANL) at 2 hours post-infection (HPI) and 24HPI for HHV-6A and HHV-6B infected cells indicate that at MOI = 1 neurites continue to extend from the time of inoculation (t_0_) through 2HPI for both HHV-6A and HHV-6B infected cells, however, by 24 HPI neurite retraction, cell death, and cell detachment from the culture surface results in insignificant neurites extension. At MOI = 2, there is a decrease in ANL between t_0_ and 2HPI with HHV-6B showing more than 35% decrease in ANL while HHV-6A results in a more modest decrease in ANL (~20%).

### 3.2.2. Neurite retraction and morphometrics (HHV-6A versus HHV-6 infected cells)

Initial manual tracing and use of a machine-learning platform (Trainabler Weka Segmentation 7) on images taken with a light microscope results in neurite and soma boundary definitions in a pseudo-fluorescence format (Fig. 6A). A pixel-by-pixel assessment by image analysis software (e.g., Neural Circuit Tracer or NeuronCytoII) produces neuronal reconstructions, from which measurements can be taken including neurite lengths and surface areas of cell soma (Fig. 6B). Morphometric analyses indicate that average neurite length in differentiated HNSCs increases between the time of inoculation (t_0_) to 2 hours post-infection (HPI) followed by a precipitous decline (retraction) between 2 HPI and 24 HPI for HHV-6A and HHV-6B infected cells at MOI=1. At MOI=2, the impact on average neurite length is more profound with a decline from t_0_ to 2 HPI in both HHV-6A and HHV-6B infected cell cultures and a precipitous decline from 2 HPI to 24 HPI. Data for MOI=2 suggest that HHV-6B infection results in greater neurite retraction than HHV-6A. For MOI=2, ~20% (N=39) and ~35% (N=14) decreases in average neurite length for HHV-6A and HHV-6B, respectively, are observed (Fig. 6C). Even at MOI=1, HHV-6B tends to attneuate neurite extension to a greater extent than HHV-6A (N=32, N=20). The magnitude of CPE severity after infection of HNSCs with virus appears to be MOI-dependent (Fig. 4, MOI = 1 versus MOI = 2). Fewer but notably long neurites extend as high-density cell clumps start to form (*see* Discussion).

### 3.3 Both glutamatergic and dopaminergic cells are susceptible to either HHV-6A or HHV-6B

To determine if either HHV-6A or HHV-6B preferentially infects select neuronal neurotransmitter phenotypes, differentiated neurons were co-labeled with neurotransmitter-specific antibodies. For glutamatergic cells, an anti-VGluT fluorescent antibody system was used to target the vesicular glutamate transporter (VGluT), a characteristic protein found in glutamatergic neurons. Simultaneous co-staining with DAPI and the anti-gB fluoroprobe demonstrate that VGluT-positive cells co-localize with the anti-gB fluorescent signal in differentiated HNSCs infected with either HHV-6A (Fig. 7, row A) or HHV-6B (Fig. 7, row B) indicating infection of glutamatergic cells. Although GFAP (a glial-specific marker) and βIII-tubulin (a neuron-specific marker) fluorescence can be detected as early as post-differentiation day (PDD) 3 with robust signal at PDD5-7, immunofluorescence signals for neurotransmitter phenotypes generally emerge from PDD7-14.

**Fig. 7.**
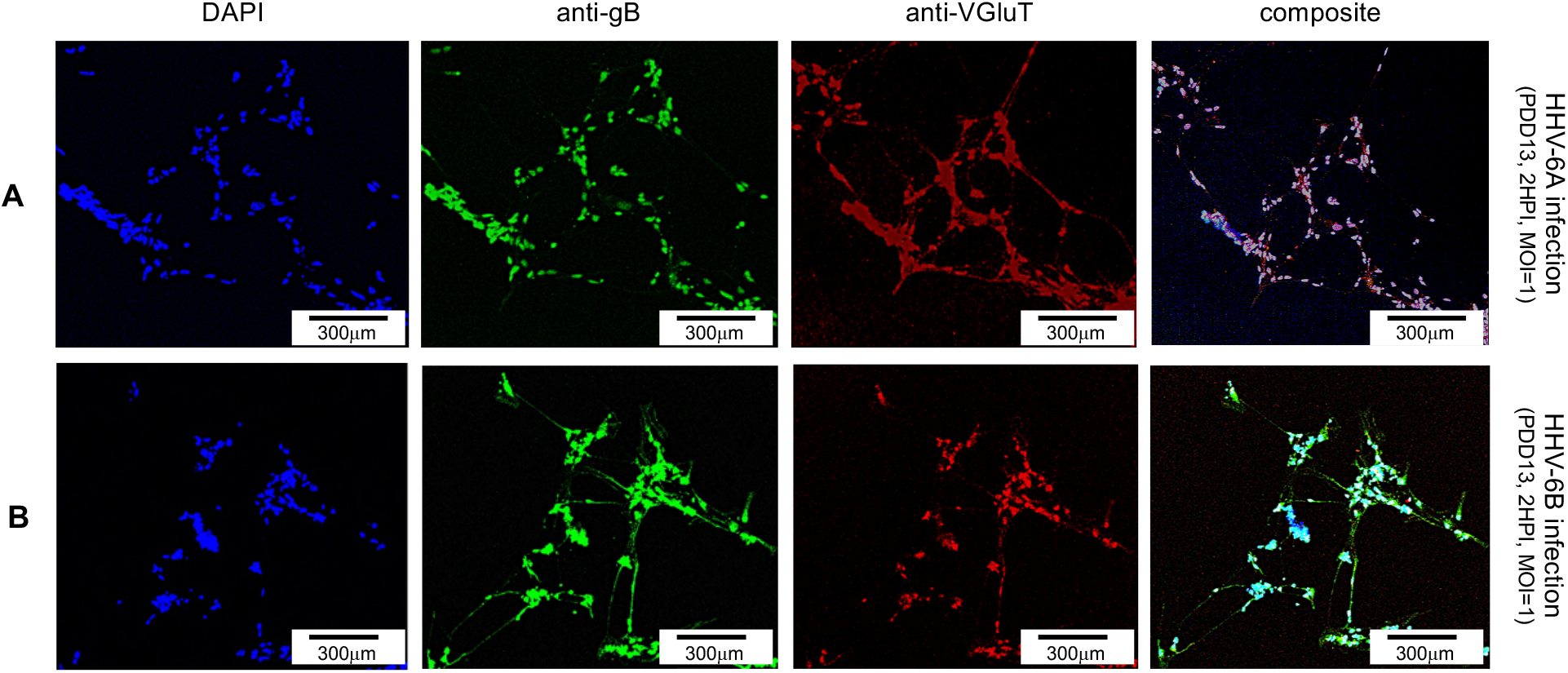
Fluorescence microscopy images of differentiated human neural stems cells (dHNSCs) treated with immunofluorescent antibodies and a fluoro-dye at PDD13: Glutamatergic Neurons. VGlut-positive dHNSCs at PDD13 (2 HPI, MOI = 1) shown (from left to right) by DAPI staining and anti-gB, and anti-VGluT immunofluorescence (with composites) indicate that gB (*green*) colocalizes with VGlut (*red*) in DAPI stained (*blue*) cells for both HHV-6A (*row A*) and HHV-6B (*row B*) infected cultures, suggesting susceptibility of glutamatergic neurons to both viruses.

For dopaminergic cells, an anti-dopamine (anti-DA) fluorescent antibody system was used to directly target dopamine. Co-staining with DAPI and co-labeling with anti-gB and anti-DA, fluorescence microscopy shows a co-localization of DAPI, anti-gB, and anti-DA signals (Fig. 8). This suggests that dopaminergic neurons are susceptible to infection by HHV-6A (Fig. 8, row A) and HHV-6B (Fig. 8, row B). In both glutamatergic and dopaminergic cells, robust HHV-6B signals for anti-gB were seen within neurites as well as cell soma.

**Fig. 8.**
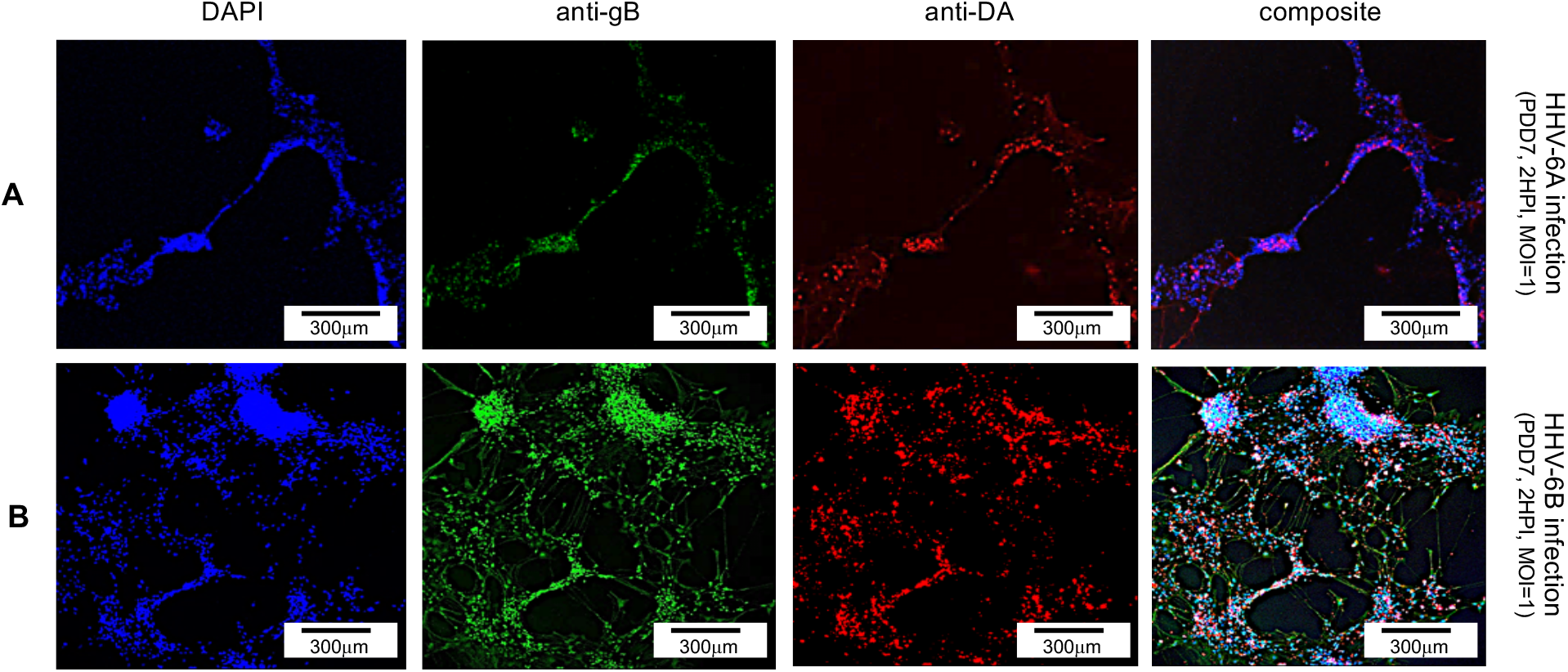
Fluorescence microscopy images of differentiated human neural stems cells (dHNSCs) treated with immunofluorescent antibodies and fluoro-dye (DAPI) at PDD7: Dopaminergic Neurons. DA-positive dHNSCs at PDD7 (2HPI, MOI = 1) shown (from left to right) by DAPI staining and anti-gB and anti-DA immunofluorescence (with composites) indicate that gB (*green*) colocalizes with dopamine (*red*) in DAPI stained (*blue*) cells for both HHV-6A (*row A*) and HHV-6B (*row B*) infected cultures, suggesting susceptibility of dopaminergic neurons to both viruses.

### 3.4 Neither HHV-6A nor HHV-6B appears to infect GAD67-positive (GABAergic) cells

To determine whether neurons that synthesize gamma-aminobutyric acid (GABA) were susceptible to infection by HHV-6A or HHV-6B, a fluorescence antibody system against GAD67, a glutamate decarboxylase that is responsible for the overwhelming majority (>90%) of GABA synthesis in the brain, was employed. (GABA is the major inhibitory neurotransmitter in the CNS).

Cultures co-stained with DAPI and co-labeled with anti-GAD67 and the anti-gB fluoroprobes failed to show infection of GAD67-positive differentiated HNSCs by either HHV-6A (Fig. 9, row A) or HHV-6B (Fig. 9, row B). Given the unexpected results, multiple time-points were examined. However, anti-gB fluorescence was not detected at PDD7 or PDD14 (Fig. 9) or under various MOIs (*data not shown*). These experiments were repeated multiple times with the same results. For one single trial (out of 5) where differentiation of HNSCs was driven toward GABA-producing cells, a few anti-gB fluorescence patches were observed in HHV-6B infected cultures (Fig. 10). Initial indications suggested that perhaps HHV-6B can indeed infected GAD67-positive cells. However, upon more detailed analysis, it appears that the colocalization of the DAPI and anti-gB signal (A1-D1 and A2-D2, respectively) does not coincide with the anti-GAD67 signal, indicating that the anti-gB fluorescence emanates from nearby cells (perhaps glia) in the mixed culture. Moreover, for some of the anti-GAD67 fluorescence there was no colocalization of a DAPI signal suggesting that these could be fluorescence signals from GABA-rich terminals. The reason for GAD67-positive cell resistance to HHV6 infection is unclear and is the subject of current research.

**Fig. 9.**
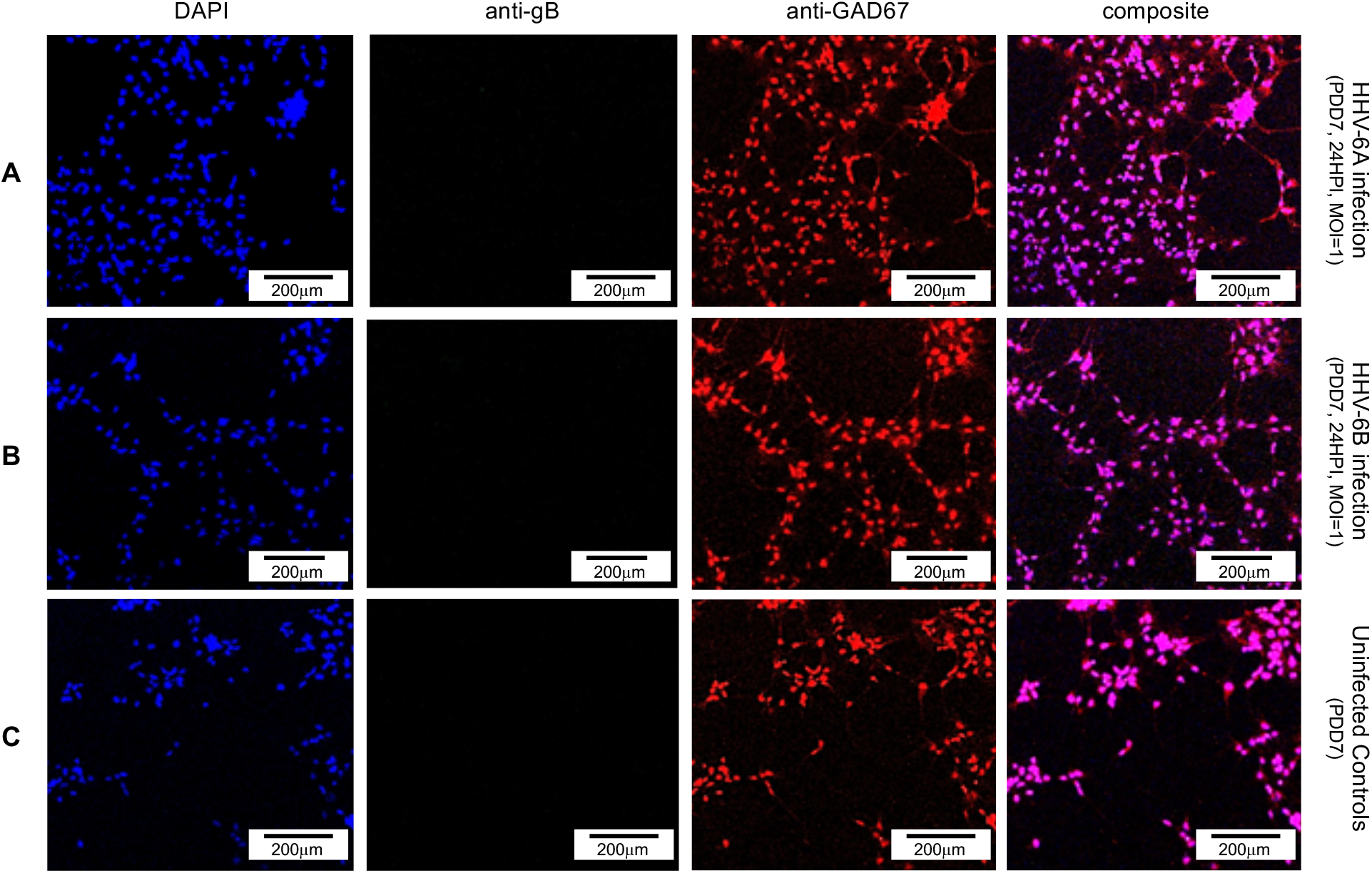
Fluorescence microscopy images of differentiated human neural stems cells (dHNSCs) treated with immunofluorescent antibodies and a fluoro-dye (DAPI) at PDD7: GABAergic Neurons. GAD67-positive dHNSCs at PDD7 (2HPI, MOI=1) shown (from left to right) by DAPI dye and anti-GAD67 immunofluorescence (with composites) indicate that gB (green) immunofluorescence does not colocalize with GAD67 (*red*) in DAPI stained (*blue*) cells for both HHV-6A (*row A*) and HHV-6B (*row B*) infected cultures, suggesting that GABAergic neurons are not susceptible to either viruses.

**Fig. 10.**
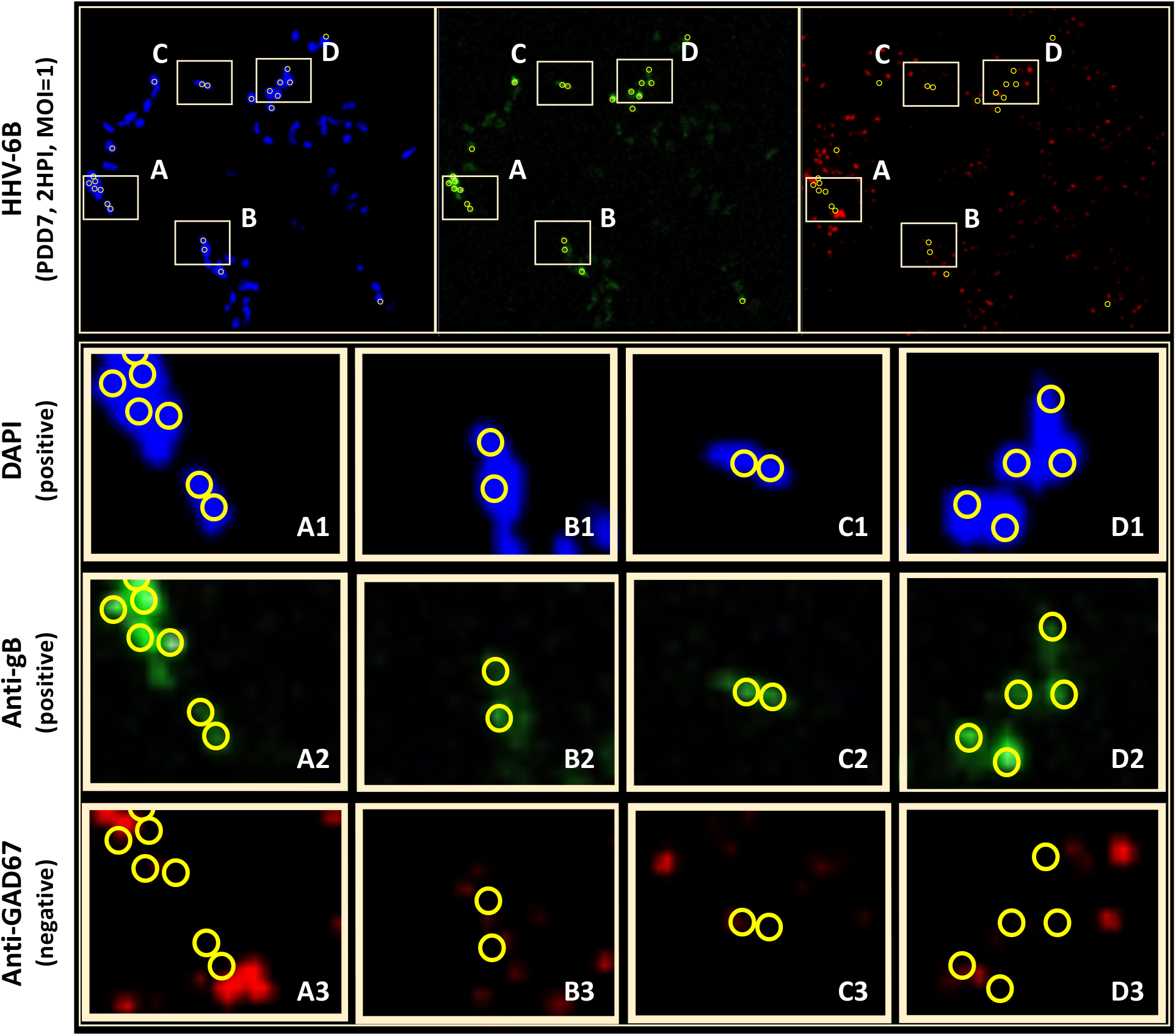
Anti-gB/DAPI fluorescence does not colocalize with GAD67-positive cells in mixed cultures. DAPI microscopy images of immunofluorescence and fluorescence staining of differentiated HNSCs at PDD7, MOI = 1, 2HPI challenged with HHV-6B. Out of multiple trials, only one anti-GAD67 positive culture displayed an anti-gB fluorescence signal indicating possible infection of GABA-containing neurons (*top*). However, upon closer examination of anti-gB cell clusters (regions A-D), it appears that DAPI positive cells (rows A1-D1) and anti-gB positive (rows A2-D2) cells are anti-GAD67 negative (row A3-D3) providing further evidence for GABAergic neuron resistance to HHV-6 infection. DAPI/anti-gB positive fluorescence is likely from adjacent glial cells or other neuronal neurotransmitter phenotypes in the mixed culture.

## 4. Discussion

Due to initial perceptions that HHV6 isolates grouped into only one virus species and the later revelation that, in fact, these are two different viruses, much of the early literature (prior to 2012) does not specify cell tropism differences between HHV-6A versus HHV-6B (Donati et al., 2003). Understanding cell tropism in the central nervous system is further confounded by the suggestion that HHV-6A may be more neurotropic than HHV-6B, however, the virus underlying viral-induced epileptogenesis (particularly, mesial temporal lobe epilepsy), is thought to be HHV-6B. Given the proposed models for HHV6-induced seizure induction, it is necessary to unravel the details of cell tropism in the CNS for these viruses. In this study, we provide novel insights and raising additional questions regarding HHV-6A versus HHV-6B cell tropism in the CNS.

### 4.1 Cytopathic effects during HHV-6 infection include syncytia formation and neurite disruption

Results from qPCR and TEM clearly indicate that productive infection of differentiated HNSCs occurs when either HHV-6A or HHV-6B is perfused into cell culture at an MOI of 0.5, 1.0, or 2.0. At the MOIs examined, results indicate that both HHV-6A and HHV-6B induce cell aggregation, syncytia formation, and cell detachment from the culture surface substrate within 30min. to 2 hrs. This suggests that there are changes in cell membrane properties during the course of infection. This is supported by morphometric analyses, which show changes in average neurite number and length as well as other morphological features (e.g., cell-cell fusion and multi-nucleation) upon infection, which are consistent with cytoskeletal breakdown or rearrangement in host cells.

Observational data over multiple trials for the MOIs tested indicate that dHNSCs infected with HHV-6B exhibit greater resistance to cell surface detachment than those infected with HHV-6A. This may be an indication that HHV-6A is more virulent on nerve cells. Cautions against generalizing data from HNSCs to *in vivo* virus-host dynamics are acknowledged. Yet, it is reasonable to suggest that these two viruses may exhibit different levels of virulence on a given host cell type. Quantitative approaches are currently ongoing in the lab to determine not only relative virulence of HHV-6A versus HHV-6B on specific cell types but also the underlying molecular substrates (i.e., transcriptomic and proteomic profiles) that can explain differences in time-course to observable cytopathic effects.

### 4.2 Susceptibility of both glial cells and neurons to HHV-6A or HHV-6B infections

Absent information about HHV-6A versus HHV-6B tropisms, prior work shows that HHV6 infects nerve cells (Albright et al., 1998; Donati et al., 2003; De Bolle et al., 2005; De Filippis et al., 2006). One study did show that HHV-6A and HHV-6B can infect astrocytes (Fotheringham et al., 2008). A separate study demonstrates HHV-6A and HHV-6B infections in neurons (Prusty et al., 2018). Yet, this study only showed HHV-6A and HHV-6B infection in Purkinje cells, which are unique. Our study also shows that both neurons (βIII tubulin-positive) and glial (GFAP-positive) cells are susceptible to HHV-6A and HHV-6B infection in cultured differentiated HNSCs (Figs. 1 and 2). However, our data also demonstrate that there is a difference in relative virulence at equivalent MOI for HHV-6A versus HHV-6B on neurons. Indeed, the time-course to the most severe CPEs is earlier for HHV-6A infection across neuronal subtypes. Specifically, HHV-6A infection induces high-density cell aggregation and cell clump detachment at early infection time-points when compared to HHV-6B infection. Although results from image analysis further demonstrate changes in neurite extension and retraction at different MOIs, these data are difficult to interpret in terms of relative virulence between viruses. This is due to the fact that neurons have been shown to extend long processes as part of a stress response during different stages of viral infection (Pettersen et al., 2006; Franze et al., 2009; Morrey et al., 2019; Kropp et al., 2020). Observational data indicated that during the course of high-density cell clump formation fewer but longer processes are extended along the culture surface until the clumps begin to detach, at which time those neurites are sheared. However, these results show that gross changes in cell morphology occur in both HHV-6A and HHV-6B infections with HHV-6A exhibiting more severe CPEs at earlier time-points post-infection.

### 4.3 Differential susceptibility of neuronal phenotypes to HHV-6A versus HHV-6B infection

Although previous work shows that neurons are susceptible to HHV6 infection, our data provide additional details regarding cell tropism for specific neuronal neurotransmitter phenotypes. Specifically, fluorescence and light microscopy show that glutamatergic (VGluT1-positive) cells and dopaminergic (DA-positive) cells are susceptible to infection by both HHV-6A and HHV-6B (Figs. 6 and 7). Although glutamatergic cells and dopaminergic cells appear to be susceptible to both viruses, infection assays failed to demonstrate that GAD67-positive (GABAergic) cells could be infected by either HHV-6A or HHV-6B. We were initially skeptical of these results since a recent study showed that Purkinje cells, which are GABAergic, are susceptible to HHV-6A and HHV-6B (Prusty et al., 2018). However, repeated trials using immunofluorescence supported by qPCR suggest that GABA neurons derived from dHNSCs are not susceptible to either virus (*see* Fig. 8). In one infection trial with HHV-6B, a sparse anti-gB fluorescence was observed in mixed cultures with GAD67-positive cells (Fig. 10, top row, middle panel). However, upon a detailed analysis, anti-GAD67 signal (*red*) does not colocalize with anti-gB/DAPI fluorescence. This suggests that HHV-6B is likely infecting cells adjacent to GABA neurons in mixed cultures and/or infected cells that are receiving GABAergic afferents. We are currently conducting experiments aimed at elucidating the molecular (i.e., -omics) underpinnings of this apparent GABA cell resistance to HHV6 infection.

### 4.4 Implications for HHV6-induced epileptogenesis

Several models are proposed as potential mechanisms by which HHV6-induced seizures can lead to epilepsy (i.e., epileptogenesis). Although neural encephalitis may emerge from primary HHV-6 infections leading to seizure (Vinters et al., 1993; Howell et al., 2012; Bartolini et al., 2019), sub-inflammatory mechanisms have also been proposed. For example, there is some *in vitro* evidence to suggest that HHV6 infection of astrocytes can lead to dysfunction in glutamate reuptake from the synapse, leading to hyperexcitation of glutamatergic pathways and thus seizure (Crawford et al., 2007; Eid et al., 2016). Beyond this *glutamate reuptake dysfunction hypothesis*, other studies have suggested that reactivation of HHV6 from latency may result in lysis of glutamatergic cells resulting in excess release of glutamate into neuronal circuits associated with seizure induction (e.g., mesial temporal lobe neural networks). This has been described as an *excitotoxicity model* (Beal et al., 1992; Bekenstein et al., 1993; Wang et al., 2007; Yao et al.,2010). If results observed in cultured HNSCs are generalizable to *in vivo* conditions, then a glutamatergic excitotoxicity model would be supported. Another hypothesis is that inhibitory interneurons which modulate glutamatergic pathways could be selectively targeted by HHV-6A or HHV-6B, thereby disrupting modulation of glutamatergic neurons, leading to excess glutamate release and seizure. If our data regarding a lack of susceptibility of GABAergic neurons to HHV6 are generalizable to *in vivo* conditions, then this latter *inhibitory (inter)neuron dysfunction hypothesis* could potentially be ruled out from the several proposed models. Of course, HHV6-induced epileptogenesis may involve more than one of these proposed models or another yet to be described mechanism. For example, cholinergic pathways from the peduncular pontine nuclei (PPN) or other tracts may also contribute to seizure induction. A lack of detail regarding: differential cell type tropisms between HHV-6A and HHV-6B; relative virulence between HHV-6A and HHV-6B on susceptible cell types; the role of HHV-7 in HHV6 reactivation; and, other unknowns motivate continued research.

## 5. Conclusions

Although the work presented is done *in vitro* using differentiated embryonic HNSCs (H-9 cells), understanding potential differences in the susceptibility of neuronal neurotransmitter phenotypes to HHV-6A versus HHV-6B allows us to more critically evaluate (and potentially, rule out) different models that seek to explain specific HHV6-induced neurological disorders (Vezzani et al., 2019). If it is determined that a given cell type is susceptible to both HHV-6A and HHV-6B, then the relative virulence and corresponding cytopathic effects (i.e., impacts and detriment to cell morphology, survival, and function) likewise should be determined to achieve a more in depth understanding of the potential role of HHV6 infections (and states of infection) in neurological disorders. This study demonstrates differential susceptibility of different neuronal neurotransmitter phenotypes as well as different time-courses of cytopathic effects (CPEs) to HHV-6A versus HHV-6B infections in dHNSCs.

## Acknowledgements

The authors thank the U.S. National Institutes of Health for providing stocks of HHV6 strains and cell lines for virus propagation. The authors also thank, Dr. Joshua Marceau for preparing working virus stocks and Elizabeth Nicole Dominguez for assisting in the manual tracing of cells as part of her undergraduate research internship. The authors thank the Arkansas Bioscience Institute for a 1-year equipment and supply grant to support this research and the U.S. National Science Foundation for supporting the work of JW via an NSF Research Experiences for Undergraduates (REU) grant (award no.1659858; PI-Ceballos).

## References

Achour, A., I. Malet, F. Le Gal, A. Dehée, A. Gautheret-Dejean, P. Bonnafous and H. Agut (2008). “Variability of gB and gH genes of human herpesvirus-6 among clinical specimens.” Journal of medical virology 80(7): 1211–1221.

Adams, M. J. and E. Carstens (2012). “Ratification vote on taxonomic proposals to the International Committee on Taxonomy of Viruses (2012).” Archives of virology 157(7): 1411–1422.

Albright, A. V., E. Lavi, J. B. Black, S. Goldberg, M. J. O’Connor and F. Gonzalez-Scarano (1998). “The effect of human herpesvirus-6 (HHV-6) on cultured human neural cells: oligodendrocytes and microglia.” Journal of neurovirology 4(5): 486–494.

Alvarez-Lafuente, R., A. Martinez, M. Garcia-Montojo, A. Mas, V. De Las Heras, M. Dominguez-Mozo, C. Maria Del Carmen, M. López-Cavanillas, M. Bartolome and E. Gomez De La Concha (2010). “MHC2TA rs4774C and HHV-6A active replication in multiple sclerosis patients.” European journal of neurology 17(1): 129–135.

Bartolini, L., W. H. Theodore, S. Jacobson and W. D. Gaillard (2019). “Infection with HHV-6 and its role in epilepsy.” Epilepsy research 153: 34–39.

Beal, M. Flint. (1992) “Mechanisms of excitotoxicity in neurologic diseases.” FASEB journal 6(15): 3338–3344.

Bekenstein, Jonathan W., and Eric W. Lothman. “Dormancy of inhibitory interneurons in a model of temporal lobe epilepsy.” Science 259, no. 5091 (1993): 97–100.

Boutolleau, D., C. Duros, P. Bonnafous, D. Caïola, A. Karras, N. De Castro, M. Ouachée, P. Narcy, M. Gueudin and H. Agut (2006). “Identification of human herpesvirus 6 variants A and B by primer-specific real-time PCR may help to revisit their respective role in pathology.” Journal of clinical virology 35(3): 257–263.

Collot, S., B. Petit, D. Bordessoule, S. Alain, M. Touati, F. Denis and S. Ranger-Rogez (2002). “Real-time PCR for quantification of human herpesvirus 6 DNA from lymph nodes and saliva.” Journal of Clinical Microbiology 40(7): 2445–2451.

Crawford, J. R., N. Kadom, M. R. Santi, B. Mariani and B. L. Lavenstein (2007). “Human herpesvirus 6 rhombencephalitis in immunocompetent children.” Journal of child neurology 22(11): 1260–1268.

De Bolle, L., J. Van Loon, E. De Clercq and L. Naesens (2005). “Quantitative analysis of human herpesvirus 6 cell tropism.” Journal of medical virology 75(1): 76–85.

De Filippis, L., C. Foglieni, S. Silva, A. Vescovi, P. Lusso and M. S. Malnati (2006). “Differentiated human neural stem cells: a new ex vivo model to study HHV-6 infection of the central nervous system.” Journal of clinical virology 37: S27–S32.

Delbrück, M. (1940). The growth of bacteriophage and lysis of the host. J. Gen. Physiol. 23: 643–660. doi: 10.1085/jgp.23.5.643

Dominguez, G., T. R. Dambaugh, F. R. Stamey, S. Dewhurst, N. Inoue and P. E. Pellett (1999). “Human herpesvirus 6B genome sequence: coding content and comparison with human herpesvirus 6A.” Journal of virology 73(10): 8040–8052.

Donati, D., N. Akhyani, A. Fogdell–Hahn, C. Cermelli, R. Cassiani-Ingoni, A. Vortmeyer, J. Heiss, P. Cogen, W. Gaillard and S. Sato (2003). “Detection of human herpesvirus-6 in mesial temporal lobe epilepsy surgical brain resections.” Neurology 61(10): 1405–1411.

Eid, T., S. E. Gruenbaum, R. Dhaher, T.-S. W. Lee, Y. Zhou and N. C. Danbolt (2016). “The glutamate– glutamine cycle in epilepsy.” The glutamate/GABA-glutamine cycle: 351–400.

Fotheringham, J., E. L. Williams, N. Akhyani and S. Jacobson (2008). “Human herpesvirus 6 (HHV-6) induces dysregulation of glutamate uptake and transporter expression in astrocytes.” Journal of Neuroimmune Pharmacology 3(2): 105–116.

Franze, K., J. Gerdelmann, M. Weick, T. Betz, S. Pawlizak, M. Lakadamyali, J. Bayer, K. Rillich, M. Gögler and Y.-B. Lu (2009). “Neurite branch retraction is caused by a threshold-dependent mechanical impact.” Biophysical journal 97(7): 1883–1890.

Hall, C. B., M. T. Caserta, K. C. Schnabel, C. Long, L. G. Epstein, R. A. Insel and S. Dewhurst (1998). “Persistence of human herpesvirus 6 according to site and variant: possible greater neurotropism of variant A.” Clinical infectious diseases 26(1): 132–137.

Howell, K. B., K. Tiedemann, G. Haeusler, M. T. Mackay, A. J. Kornberg, J. L. Freeman and A. S. Harvey (2012). “Symptomatic generalized epilepsy after HHV6 posttransplant acute limbic encephalitis in children.” Epilepsia 53(7): e122–e126.

Isegawa, Y., T. Mukai, K. Nakano, M. Kagawa, J. Chen, Y. Mori, T. Sunagawa, K. Kawanishi, J. Sashihara and A. Hata (1999). “Comparison of the complete DNA sequences of human herpesvirus 6 variants A and B.” Journal of virology 73(10): 8053–8063.

Kropp, K. A., A. D. López-Muñoz, B. Ritter, R. Martín, A. Rastrojo, S. Srivaratharajan, K. Döhner, A. Dhingra, J. S. Czechowicz and C.-H. Nagel (2020). “Herpes simplex virus 2 counteracts neurite outgrowth repulsion during infection in a nerve growth factor-dependent manner.” Journal of Virology 94(20).

Liu, D., X. Wang, Y. Wang, P. Wang, D. Fan, S. Chen, Y. Guan, T. Li, J. An and G. Luan (2018). “Detection of EBV and HHV6 in the Brain Tissue of Patients with Rasmussen’s Encephalitis.” Virologica Sinica 33(5): 402–409.

Michael, B. D. and T. Solomon (2012). “Seizures and encephalitis: clinical features, management, and potential pathophysiologic mechanisms.” Epilepsia 53: 63–71.

Mohammadpour Touserkani, F., M. Gaínza-Lein, S. Jafarpour, K. Brinegar, K. Kapur and T. Loddenkemper (2017). “HHV-6 and seizure: A systematic review and meta-analysis.” Journal of medical virology 89(1): 161–169.

Morrey, J. D., A. L. Oliveira, H. Wang, K. Zukor, M. V. de Castro and V. Siddharthan (2019). “Zika virus infection causes temporary paralysis in adult mice with motor neuron synaptic retraction and evidence for proximal peripheral neuropathy.” Scientific reports 9(1): 1–15.

Oyaizu, H., H. Tang, M. Ota, N. Takenaka, K. Ozono, K. Yamanishi and Y. Mori (2012). “Complementation of the function of glycoprotein H of human herpesvirus 6 variant A by glycoprotein H of variant B in the virus life cycle.” Journal of virology 86(16): 8492–8498.

Pettersen, J. A., G. Jones, C. Worthington, H. B. Krentz, O. T. Keppler, A. Hoke, M. J. Gill and C. Power (2006). “Sensory neuropathy in human immunodeficiency virus/acquired immunodeficiency syndrome patients: Protease inhibitor–mediated neurotoxicity.” Annals of Neurology: Official Journal of the American Neurological Association and the Child Neurology Society 59(5): 816–824.

Prusty, B. K., N. Gulve, S. Govind, G. R. Krueger, J. Feichtinger, L. Larcombe, R. Aspinall, D. V. Ablashi and C. T. Toro (2018). “Active HHV-6 infection of cerebellar purkinje cells in mood disorders.” Frontiers in microbiology 9: 1955.

Rapp, J. C., L. T. Krug, N. Inoue, T. R. Dambaugh and P. E. Pellett (2000). “U94, the human herpesvirus 6 homolog of the parvovirus nonstructural gene, is highly conserved among isolates and is expressed at low mRNA levels as a spliced transcript.” Virology 268(2): 504–516.

Raspall-Chaure, M., T. Armangué, I. Elorza, À. Sanchez-Montanez, M. Vicente-Rasoamalala and A. Macaya (2013). “Epileptic encephalopathy after HHV6 post-transplant acute limbic encephalitis in children: confirmation of a new epilepsy syndrome.” Epilepsy research 105(3): 419–422.

Santoro, F., H. L. Greenstone, A. Insinga, M. K. Liszewski, J. P. Atkinson, P. Lusso and E. A. Berger (2003). “Interaction of glycoprotein H of human herpesvirus 6 with the cellular receptor CD46.” Journal of Biological Chemistry 278(28): 25964–25969.

Santoro, F., P. E. Kennedy, G. Locatelli, M. S. Malnati, E. A. Berger and P. Lusso (1999). “CD46 is a cellular receptor for human herpesvirus 6.” Cell 99(7): 817–827.

Santpere, G., M. Telford, P. Andrés-Benito, A. Navarro and I. Ferrer (2020). “The Presence of Human Herpesvirus 6 in the Brain in Health and Disease.” Biomolecules 10(11): 1520.

Tang, H. and Y. Mori (2018). Glycoproteins of HHV-6A and HHV-6B. Human Herpesviruses, Springer: 145–165.

Tang, H., S. Serada, A. Kawabata, M. Ota, E. Hayashi, T. Naka, K. Yamanishi and Y. Mori (2013). “CD134 is a cellular receptor specific for human herpesvirus-6B entry.” Proceedings of the National Academy of Sciences 110(22): 9096–9099.

Vezzani, A., S. Balosso and T. Ravizza (2019). “Neuroinflammatory pathways as treatment targets and biomarkers in epilepsy.” Nature Reviews Neurology 15(8): 459–472.

Vinters, H. V., R. Wang and C. A. Wiley (1993). “Herpesviruses in chronic encephalitis associated with intractable childhood epilepsy.” Human pathology 24(8): 871–879.

Wang, Fu-Zhang, and Philip E. Pellett. “HHV-6A, 6B, and 7: immunobiology and host response.” Human herpesviruses: biology, therapy, and immunoprophylaxis (2007).

Yao, K., J. R. Crawford, A. L. Komaroff, D. V. Ablashi and S. Jacobson (2010). “Review part 2: Human herpesvirus-6 in central nervous system diseases.” Journal of medical virology 82(10): 1669

